# *N*^6^-methyladenosine promotes temozolomide resistance through mRNA stabilization in glioblastoma cells

**DOI:** 10.1101/2025.09.14.676126

**Authors:** Emily Dangelmaier, Dorthy Fang, Kyal Sin Htet, Nicole S. Harry, Renee Pascoe, Je Won Yang, Clara Wang, Sigrid Nachtergaele

## Abstract

*N*^6^-methyladenosine (m^6^A) is a critical regulator of mRNA processing and function, impacting nearly every step of the mRNA lifecycle. Changes in m^6^A status have been implicated in many different types of cancer, including glioblastoma where acquired resistance to temozolomide remains a major clinical challenge. We established glioblastoma cell culture models of acquired temozolomide resistance and analyzed the role of m^6^A in controlling resistance-associated pathways. We show that m^6^A stabilizes key genes and pathways promoting temozolomide resistance in glioblastoma, including *MGMT*, the enzyme that repairs primary temzolomide-induced DNA damage, and PI3K/Akt signaling, a major driver of chemoresistance. Pharmacological inhibition of the m^6^A methyltransferase METTL3 destabilizes *MGMT* and other critical mRNAs, restoring temozolomide sensitivity. Collectively, our results suggest that genes associated with temozolomide resistance are stabilized by m^6^A methylation, even as the majority of the transcriptome remains subject to m^6^A-mediated mRNA decay. Moreover, these data highlight METTL3 inhibition as a promising therapeutic approach to overcoming temozolomide resistance in glioblastoma.

## INTRODUCTION

Chemical modifications are critical regulators of the structure and function of diverse RNA species [1,2]. Recent work has highlighted the importance of modifications in mRNA processing and function, revealing their roles in mRNA capping, splicing, polyadenylation, export, translation, and decay [3–5]. *N*^6^-methyladenosine (m^6^A) is the most prevalent internal mRNA modification, occurring at approximately 0.5% of adenosines [6–8]. Transcriptome-wide mapping of m^6^A sites [6,9] and identification of m^6^A regulatory machinery [5,7,10–18] have spurred rapid progress in our understanding of the m^6^A function. For example, m^6^A can facilitate mRNA decay via the binding of YTHDF2 and recruitment of deadenylation and decay machinery [12,13], and this mechanism has regulatory roles in many biological contexts including embryonic development [19–21], stem cell differentiation [22,23], and cancer [24–27].

As additional m^6^A-binding proteins and regulatory mechanisms have been discovered, a more complicated picture of m^6^A-mediated gene expression regulation has emerged [5,10]. For example, the IGF2BP proteins have been reported to stabilize specific mRNAs in an m^6^A-dependent manner [16], but how or why a specific m^6^A site is bound by one binding protein versus another remains an open question. Recent work has also suggested that direct interactions between m^6^A and the ribosome can cause ribosome stalling, inducing mRNA decay via non-YTHDF2 mediated mechanisms [28–30].

Adding another layer of complexity, m^6^A methylation can also alter RNA structure, which can occlude or expose regulatory sequences depending on local RNA folding interactions [31,32]. All of these mechanisms of m^6^A action – specific interactions with binding proteins, interactions with translation and other cellular machinery, changes in RNA structure, and perhaps others not yet identified – could be operating simultaneously in cells to shape the transcriptome. However, how these mechanisms coexist and how groups of transcripts might be differentially regulated by different mechanisms within the same cells remains unclear. Moreover, a downstream consequence of these multiple m^6^A regulatory modes is the fact that the functional outcomes of m^6^A methylation on a given mRNA can change depending on the specific gene expression profile of the cell [33].

Given such wide ranging impacts of m^6^A on gene expression regulation, it is not surprising that changes in mRNA m^6^A status and in expression levels of m^6^A-associated regulatory factors have been implicated in many human cancers [1,34]. A common theme is that changes in m^6^A occupancy on key oncogene or tumor suppressor transcripts regulates their expression, which in turn influences tumor progression, treatment response, and overall patient survival. It is also clear, however, that m^6^A studies focused on the same disease can come to different conclusions as to the tumor suppressive or oncogenic effects of m^6^A in that context, which could be a consequence of different combinations of mechanisms highlighted above [35]. Nevertheless, the role of m^6^A in oncogenic transformation motivated the development of small molecule inhibitors of the m^6^A mRNA methyltransferase METTL3, such as STM2457 [36]. STC-15, an orally bioavailable derivative of STM2457 is in clinical trials for a variety of solid tumors [37]. METTL3 inhibitors have also been explored in the context of adjuvant therapy to enhance the effects of current standard of care chemotherapies [38–41].

Acquired resistance to frontline cancer therapies is a significant challenge in the clinic, and remains a particularly formidable hurdle in the treatment of glioblastoma (GBM) [42,43]. After surgical resection, patients are treated with radiation therapy and the alkylating agent temozolomide (TMZ) [44]. While many patients respond initially, the overall 5-year survival rate is estimated to be less than 10%, in large part due to frequent tumor recurrence, initial or acquired resistance to TMZ, and limited alternative treatment options [43]. O^6^-methylguanine-DNA methyltransferase (MGMT) is an enzyme that repairs cytotoxic DNA lesions caused by TMZ, reversing TMZ-induced DNA damage. DNA methylation of the *MGMT* promoter silences its expression and is associated with improved response to TMZ in GBM patients [45,46]. While efforts have been made to sensitize GBM to TMZ with MGMT small molecule inhibitors, these drugs demonstrated high toxicity in early clinical trials [47].

Despite rapid progress in our understanding of RNA modifications in cancer, including GBM [26,48,49], less is known about their roles in drug resistance. Given the wide ranging impacts of m^6^A methylation on gene expression programs in cancer and the significant challenges confronting GBM patients in the clinic, we were motivated to explore the functions of mRNA m^6^A methylation in TMZ resistance. We hypothesized that the large-scale cellular changes that occur during acquired drug resistance could be regulated by m^6^A, and that better understanding these processes could expose much needed novel therapeutic strategies to combat resistance.

Here, we develop cell culture models of acquired TMZ resistance in GBM to investigate whether key resistance-associated genes and pathways are directly regulated by m^6^A. In GBM cell lines, prolonged TMZ exposure induced durable resistance accompanied by massive upregulation of MGMT expression, likely driven by demethylation of its promoter. Treatment with the METTL3 inhibitor STM2457 is lethal to both parental and TMZ-resistant cell lines, consistent with what has been observed in other tumors. We also find that *MGMT* mRNA is m^6^A-methylated and remarkably stable, helping to sustain high MGMT expression in TMZ-resistant cells. STM2457 treatment destabilizes *MGMT* mRNA, reducing its expression and re-sensitizing cells to TMZ. We also find that other genes that are upregulated in our TMZ-resistant cells tend to be regulated in a similar manner, suggesting that m^6^A-mediated mRNA stabilization of TMZ-resistance associated genes may represent a broader mechanism that helps to maintain stable TMZ resistance. Taken together, our findings provide a molecular foundation supporting the possibility that m^6^A inhibition may be a viable therapeutic avenue for drug-resistant GBM.

## RESULTS

### Generation of TMZ-resistant cultured glioma cell lines

To explore whether RNA modifications regulate TMZ resistance, we generated paired TMZ-sensitive and TMZ-resistant cultured glioma cells. We chose the parental cell lines based on 1) ease of manipulation and scalability in culture, and 2) low basal *MGMT* expression, an indicator of baseline sensitivity to TMZ [45]. We chose U87 MG, LN229, and U251 MG cell lines as they exhibit both properties (Figure 1A) [50]. In addition, we used T98G cells as a control that has high basal *MGMT* expression and is resistant to TMZ. To generate our paired TMZ-sensitive and TMZ-resistant (TMZ-R) cell lines, we continually cultured U87 MG, LN229, and U251 MG cells with 50 μM TMZ (Figure 1B). Following an initial phase of high cell death, a small subpopulation of cells survived and achieved stable TMZ-resistance after approximately four weeks. Thereafter, these resistant cells could be continually cultured in the presence of TMZ. We refer to these TMZ-resistant cell lines as U87-R, LN229-R, and U251-R, and their parental TMZ-sensitive counterparts by their standard names. To test for stable TMZ resistance, we assessed the effect of acute TMZ treatment on cell viability, clonogenicity, and cell growth of these cells and their TMZ-sensitive counterparts. While parental U87 MG, LN229, and U251 MG cells showed reduced viability, clonogenicity, and growth upon TMZ treatment, U87-R, LN229-R, and U251-R cells were unaffected (Figures 1C-D, 2D).

**Figure 1.**
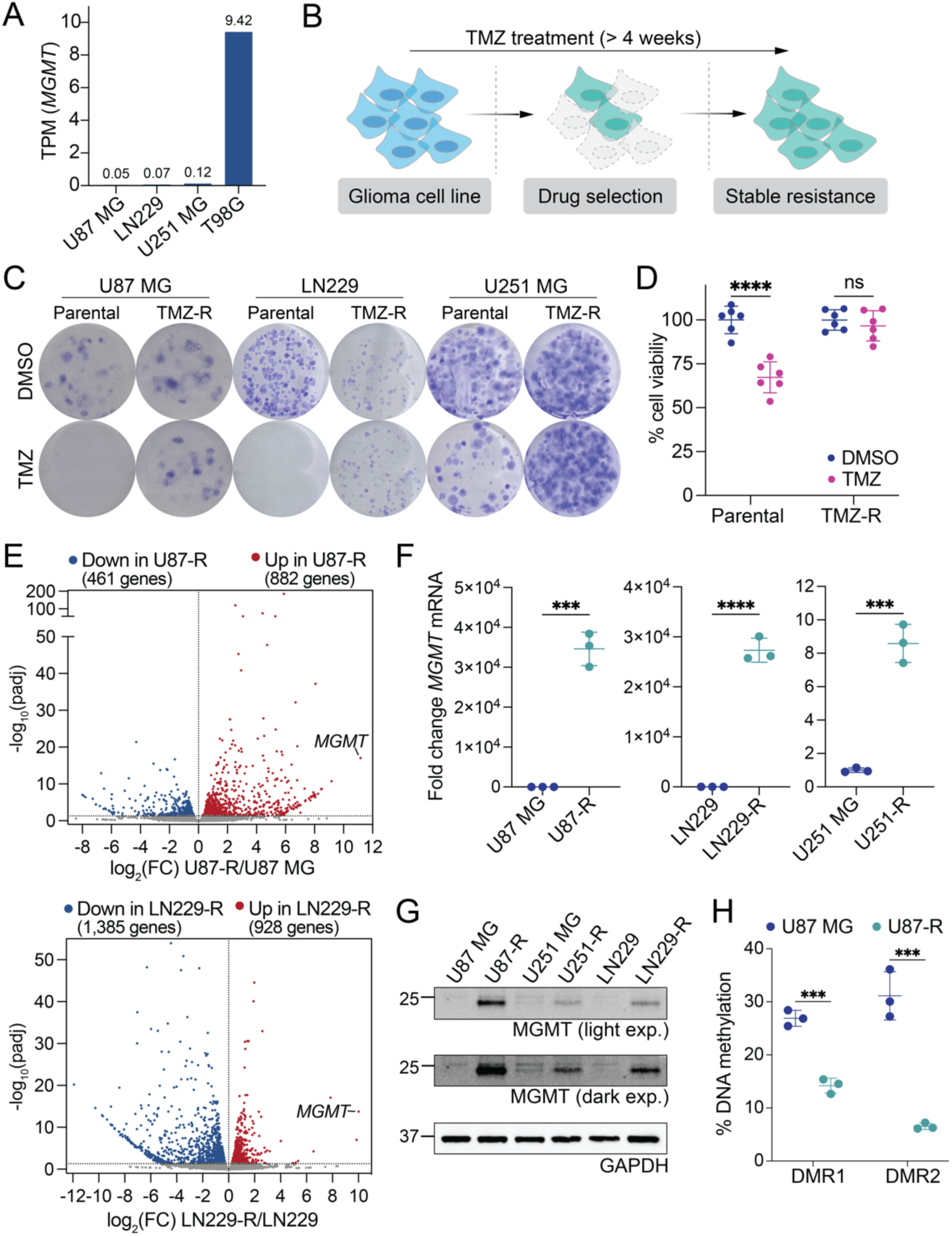
Modelling acquired TMZ resistance in GBM. (A) Cancer Cell Line Encyclopedia (CCLE [50]) RNA-seq gene expression data for *MGMT* (transcripts per million) in cell lines used in this study. (B) Strategy for generating TMZ-resistant (TMZ-R) cell lines. Cells were continuously maintained with 50 μM TMZ for >4 weeks. (C) Representative images of colony formation assays for parental and TMZ-R cells treated with 50 μM TMZ or DMSO. (D) Treatment with 50 μM TMZ for 96h reduced cell viability of U87 MG parental, but not TMZ-R, cells (*n* = 6). (E) Volcano plots showing differentially expressed genes in U87-R or LN229-R compared to respective parental lines. Genes with adjusted p-values (padj) < 0.05 are considered differentially expressed. (F) *MGMT* mRNA fold change in TMZ-R cells relative to parental lines using *GAPDH* as an internal control (*n* = 3). (G) Western blot analysis for MGMT in parental and TMZ-R cells. A light and dark exposure of the same blot are shown to more easily observe weaker signals. GAPDH served as a loading control. (H) Quantification of DNA methylation level at differentially methylated region (DMR) 1 and 2 of the *MGMT* promoter CpG island in U87 MG parental and TMZ-resistance cells. Data is mean ± s.d. of 3 biological replicates. Data are mean ± standard deviation (s.d.). Two-tailed Student’s *t*-test; ***p < 0.001; ****p < 0.0001; ns, not significant.

To interrogate mechanisms driving TMZ resistance, we performed RNA-seq in U87 MG, U87-R, LN229, and LN229-R cells to profile TMZ resistance-related gene expression changes transcriptome-wide (Figure 1E, Table S1). Overall, genes differentially expressed (DEGs) between TMZ-R and parental cells were enriched for KEGG pathways known to be associated with TMZ resistance, including PI3K-Akt, MAPK, and P53 signaling pathways, as well as extracellular matrix-receptor interactions (Figure S1A) [51–53]. DEGs were also significantly enriched for genes associated with TMZ in the Comparative Toxicogenomic Database (CTD) (Figure S1B) [54].

Although U87 MG and LN229 cells have distinct gene expression profiles, in both cases the most highly upregulated transcript in the respective TMZ-R cell lines is the GBM prognostic marker *MGMT* (Figure 1E). We validated these results in additional biological replicates from the U87 MG and LN229 lines and in U251 MG cells using quantitative PCR (RT-qPCR) (Figure 1F). Critically, we also observe a correlated increase in MGMT protein by immunoblotting (Figure 1G), suggestive of upregulated MGMT enzyme activity in the TMZ-R cells. This substantial MGMT upregulation reflects what is seen clinically in GBM patients who are resistant to TMZ, because high MGMT activity is critical for repairing TMZ-induced DNA damage [45,46]. We also observed that two genomic regions whose methylation status is known to influence *MGMT* transcriptional activation in patients, demethylated regions 1 and 2 (DMR1 and DMR2) [55], were also demethylated in our U87-R cells relative to U87 MG cells (Figure 1H). Taken together, these data, particularly the upregulation of MGMT mediated via promoter demethylation, gives us confidence that we can use our TMZ-R cells as a cellular model to reveal molecular mechanisms that regulate TMZ resistance.

### TMZ-resistant cell lines are sensitive to METTL3 inhibition

METTL3-mediated m^6^A methylation of mRNA has been implicated in a variety of cancers (reviewed in [1]), spurring development of small molecule inhibitors of METTL3 such as STM2457 [36]. Inhibition of METTL3 can also improve sensitivity to other therapies [38–41], and previous work has shown that loss of METTL3 function can be lethal [22,23]. We were therefore curious to see how METTL3 inhibition and loss of m^6^A would impact our glioma cell lines, particularly the TMZ-R lines. To test STM2457 efficacy and tune dosing for subsequent experiments, we treated U87 MG cells with 10 μM and 50 μM STM2457 for 72 hours. We analyzed bulk level m^6^A changes in total RNA and mRNA by liquid chromatography coupled to tandem mass spectrometry (LC-MS/MS). To isolate highly purified mRNA, we performed poly(A) selection, ribosomal RNA (rRNA) depletion, and then size selection to remove small RNAs (Figure 2A); we validated successful purification by automated electrophoresis. Both the 10 μM and 50 μM doses of STM2457 resulted in a significant reduction in m^6^A level in mRNA (Figure 2A) with minimal effects on cell viability in this time frame (Figure 2B). In contrast, METTL3 inhibition does not affect the m^6^A level in total RNA, which contains m^6^A from abundant rRNAs that is installed by other methyltransferases [56–59]. This is expected given the small fraction of total RNA that mRNA represents, and is consistent with the previously reported specificity of this inhibitor for the mRNA m6A methyltransferase METTL3 [36].

**Figure 2.**
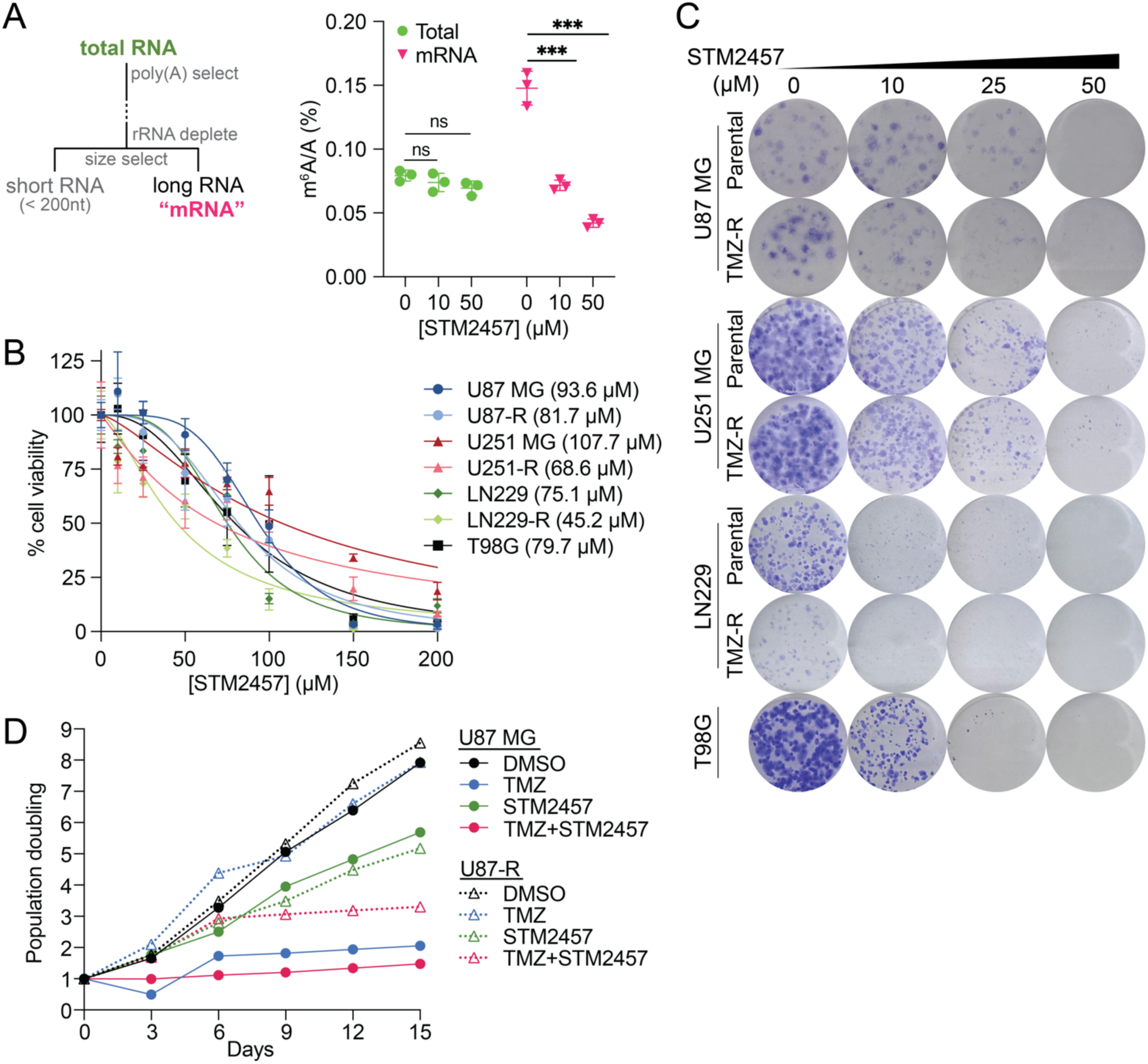
METTL3 inhibition reduces cell growth and improves TMZ response in glioma cells. (A) LC-MS/MS quantification of m^6^A levels expressed as percent m^6^A over unmodified adenosine in total RNA and mRNA from U87 MG cells treated with STM2457 for 72 h. mRNA was purified by poly(A) selection, rRNA depletion, and size selection (>200 nt). Data are mean ± s.d, *n* = 3 injections (***p < 0.001, ns, not significant, two-tailed Student’s *t*-test). (B) Dose-response curves for glioma cell lines treated with increasing concentrations of STM2457 for 96 h. IC50 for each cell line is shown in parentheses. Data are mean ± s.d., *n* = 6. (C) Representative images of colony formation assays of glioma cells treated with STM2457 (P = Parental; R = TMZ-resistant). (D) Population doublings in U87 MG and U87-R cells treated with DMSO alone, 50 μM TMZ, 50 μM STM2457, or a combined dose of 50 μM TMZ and 50 μM STM2457.

Because METTL3 inhibition has been previously shown to have anti-cancer effects [36,40,60,61], we further investigated the effects of different STM2457 doses and treatment times on our glioma cell lines. We also included T98G cells, which express high levels of *MGMT* (Figure 1A) and are resistant to TMZ at baseline. Despite their dramatically different responses to TMZ (Figure 1C-D, 2D), STM2457 treatment reduces cell viability, clonogenicity, and growth in both the parental and TMZ-R lines (Figure 2B-D). We also noted a small but consistent increase in sensitivity to STM2457 in the TMZ-R lines relative to their TMZ-sensitive counterparts, indicated by the respective IC50s in DNA-dye based cell viability assays (Figure 2B). Cell viability in the presence of STM2457 highly depends not only on the dose, but also on treatment time and density of cells at the time of treatment. The sparse cell density and 14-day treatment time used for colony formation assays increased sensitivity to STM2457, and under these conditions the 50 μM dose severely inhibited colony formation in parental and TMZ-R cells (Figure 2C). We also measured the ability of cells to grow over multiple passages in TMZ, STM2457, or both drugs together. Though U87-R cells are insensitive to the presence of TMZ alone, their doubling rate drops in the presence of STM2457 (Figure 2D). This effect is even more pronounced when TMZ is combined with STM2457 (Figure 2D), suggesting that METTL3 inhibition can resensitize cells to TMZ.

### Bulk level m^6^A does not change with chronic or acute TMZ treatment

Inspired by observations in other cancers, we suspected that critical transcripts in our glioma cells are m^6^A-methylated [26,35,48,49]. We first measured bulk levels of m^6^A in parental and TMZ-R cells, and did not detect significant differences in m^6^A levels in total RNA or mRNA between U87 MG and U87-R cells, nor the levels of METTL3 and its obligate binding partner METTL14 (Figure S2A-C). We next wondered whether acute TMZ treatment could result in bulk level m^6^A changes, and performed LC-MS/MS on RNA from U87 MG cells acutely treated with 100 μM TMZ for 72 hours.

To get a more detailed picture of possible changes in different RNA sub-populations, we used a more comprehensive series of purifications that resulted in five different subpopulations of RNA: total RNA, short RNA (tRNA and small RNA species shorter than 200 nucleotides), long RNA (mRNA, rRNA, and lncRNA greater than 200 nucleotides long), long/polyA+ RNA (enriched for polyadenylated RNAs including mRNA, but with possible remaining rRNA contamination), and long/rRNA- RNA (depleted for rRNA species, enriched for mRNAs and lncRNAs regardless of polyA status) (Figure S2D). Once again, there were minimal detectable differences, although we did observe a small but statistically significant increase in m^6^A levels in the long/rRNA- fraction, perhaps suggesting that acute TMZ treatment may result in elevated m^6^A methylation of some mRNA or lncRNA transcripts (Figure S2D).

### METTL3 inhibition alters expression of key regulatory pathways in glioma cells

We next considered whether a subset of transcripts critical to glioma cell growth and TMZ resistance are affected by METTL3 inhibition. We performed RNA-seq in U87 MG, U87-R, LN229, and LN229-R cells treated with or without 50 μM STM2457 for 96 hours (Table S2, Figure 2A-B). Overall, the majority of DEGs were upregulated in response to STM2457 treatment (Figure 3A, Figure S3A), consistent with previous observations and the established role of m^6^A in destabilizing mRNA [12,13]. KEGG pathway and Gene Ontology Biological Pathway (GOBP) analysis revealed that genes upregulated by STM2457 in TMZ-R cells were enriched for pathways related to autophagy, apoptosis, cell death and catabolism, whereas downregulated genes were enriched for pathways related to cell division and development (Figure 3B, Figure S3B-C). Genes downregulated by STM2457 treatment were also enriched for pathways important for promoting TMZ resistance, including DNA repair pathways, hypoxia response/HIF-1 signaling, PI3K-Akt signaling, TNF signaling, and glycolysis. This observation is consistent with our findings that STM2457 treatment decreased growth of glioma cells and improved TMZ sensitivity of resistant cells (Figure 2B-D).

**Figure 3.**
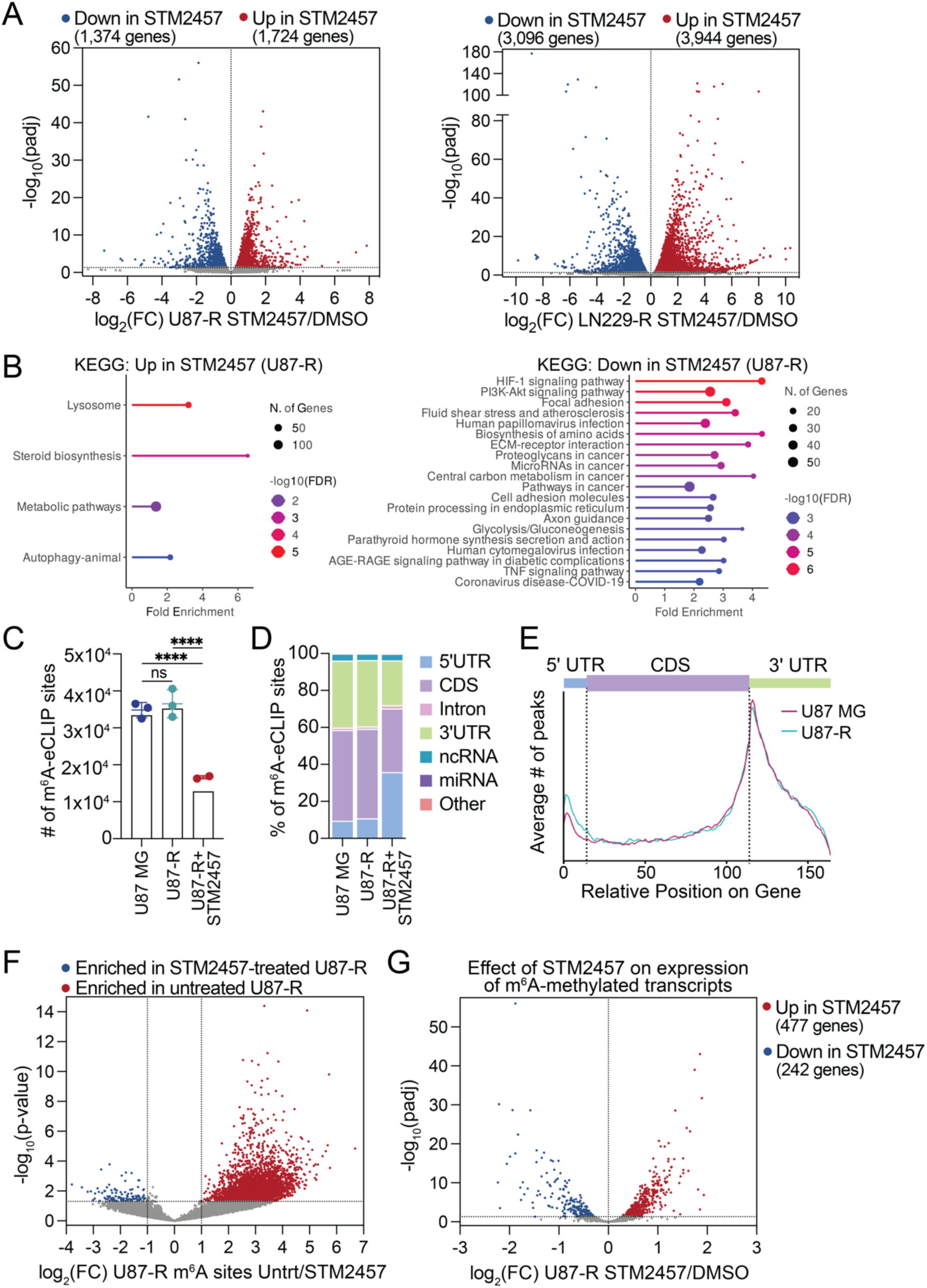
Transcriptome-wide regulation by m6A in TMZ-resistant GBM. (A) Volcano plots showing differentially expressed genes (padj < 0.05) in U87-R or LN229-R treated with 50 μM STM2457 for 96 h. (B) KEGG pathways enriched for genes differentially expressed in response to STM2457 treatment in U87-R cells. (C) Total number of single-nucleotide m^6^A-eCLIP sites. Bars represent the number of reproducible sites in each condition while points represent values for individual replicates with mean ± s.d. (****p < 0.0001; ns, not significant; two-tailed Student’s *t*-test). (D and E) Genomic distribution of m^6^A. (D) Percent of reproducible m^6^A sites in each condition at each feature. (E) Representative metagene analysis showing distribution of reproducible m^6^A-eCLIP peaks in U87 MG parental (magenta) and U87-R (turquoise). (F) Volcano plot showing differential enrichment of m^6^A-eCLIP sites in U87-R cells treated with STM2457 compared to untreated. (G) Volcano plot showing the effect of STM2457 treatment on the expression of transcripts which have STM2457-sensitive m^6^A sites (reduced enrichment of log2 fold change ≥ 1; p-value < 0.05) in U87-R cells (transcripts in red in 3F).

To gain further insight into the direct mechanisms by which m^6^A regulates GBM tumorigenesis and TMZ response, we performed m^6^A enhanced crosslinking and immunoprecipitation (m^6^A-eCLIP) in U87 MG, U87-R, and U87-R treated with STM2457 to reveal transcript- and site-specific differences in m^6^A status (Table S3). In brief, RNA was fragmented and incubated with anti-m^6^A antibody before being crosslinked by UV light, and antibody-bound RNAs were then identified by high throughput sequencing [62,63]. Using this method, we were able to map m^6^A in the transcriptome at high resolution. To increase the rigor of our experiment, we used a sub-lethal STM2457 treatment as a specificity control to help filter out non-METTL3-mediated m^6^A sites in these anti-m^6^A immunoprecipitations.

Consistent with our LC-MS/MS findings that bulk m^6^A levels are similar between U87 MG and U87-R cells (Figure S2A), m^6^A-eCLIP revealed no significant difference in the number of called m^6^A sites between the two cell lines (Figure 3C). Additionally, there were no discernible differences in m^6^A deposition patterns between U87 MG and U87-R cells. In both cell lines, the majority of m^6^A sites were found at canonical DRACH motifs (where D = A, G, or U; R = A or G; and H = A, C, or U) located at the end of coding sequences and early in 3′UTRs near stop codons (Figure 3D-E, Figure S3D), which is typical of METTL3-mediated m^6^A methylation [6,9]. STM2457 treatment significantly reduced the number and enrichment of m^6^A sites (Figure 3C,F, Table S4). Of the m^6^A sites that remain in STM2457-treated U87-R cells, a smaller proportion are in coding sequences, 3′UTRs, and at DRACH motifs compared to untreated U87-R cells (Figure 3D-E, Figure S3D). Intersecting our m^6^A-eCLIP and RNA-seq datasets, we defined a set of high confidence m^6^A sites based on reduced enrichment in STM2457 treated samples relative to controls (Figure 3F, transcripts in red), and examined the effect of METTL3 inhibition on their expression level (Figure 3G). m^6^A-methylated transcripts were both up- and downregulated by METTL3 inhibition, though the majority of transcripts were upregulated, as expected (Figure 3G). Taken together, these results validate that STM2457 treatment effectively disrupts METTL3-mediated m^6^A deposition in glioma cells, and reveals that key cellular processes central to TMZ resistance are regulated by m^6^A.

### *MGMT* mRNA is m^6^A-methylated and stabilized via binding of IGF2BP2

We next investigated whether m^6^A plays a direct role in TMZ resistance. We examined m^6^A-eCLIP single nucleotide sites which were differentially enriched between U87 MG and U87-R cells (Table S4). Strikingly, two of the most highly significantly enriched m^6^A sites in U87-R cells were located on *MGMT* (Figure 4A). At the transcript level, *MGMT* was also significantly enriched in U87-R m^6^A-eCLIP but not in STM2457-treated U87-R m^6^A-eCLIP (Figure 4B). Both called m^6^A single-nucleotide sites lie within an m^6^A-eCLIP enrichment peak, are at DRACH motifs, and are positioned within the coding sequence proximal to the stop codon. To further validate *MGMT* m^6^A-methylation, we immunoprecipitated (IP) polyA-enriched RNA isolated from U87-R, LN229-R, and T98G cells with an anti-m^6^A antibody and quantified enrichment by RT-qPCR. *MGMT* mRNA was significantly and reproducibly enriched in the anti-m^6^A IP samples in all cell lines, at a level similar to that of other known m^6^A-methylated RNAs such as *MALAT1* (Figure 4C, Figure S4A-D). STM2457 treatment resulted in no significant enrichment of *MGMT* mRNA by m^6^A-IP (Figure S4D).

**Figure 4.**
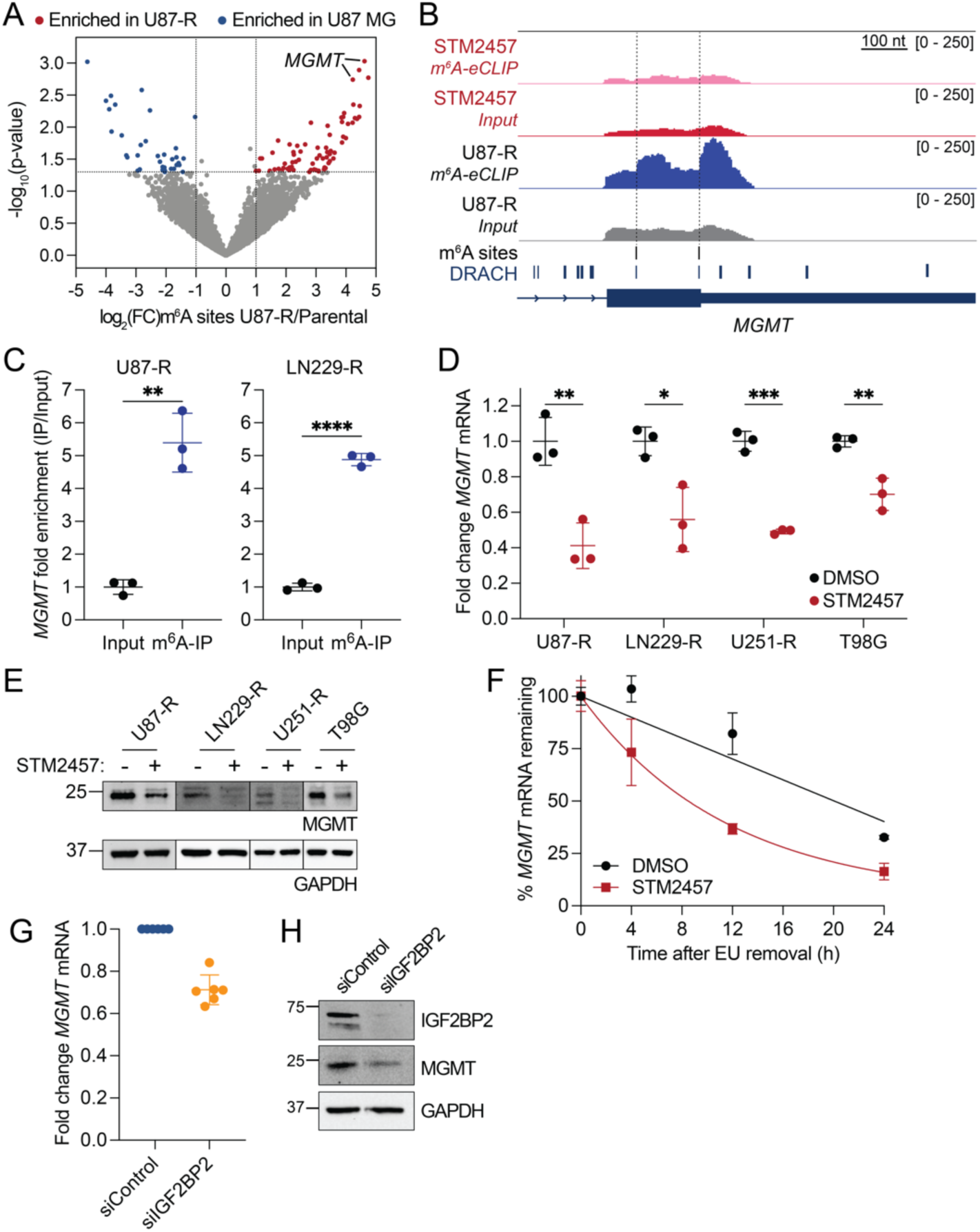
*MGMT* is post-transcriptionally regulated by m^6^A. (A) Volcano plot of m^6^A-eCLIP sites differentially enriched in U87-R cells compared to U87 MG. m^6^A sites in *MGMT* are labeled. (B) Integrative Genomics Viewer (IGV) snapshot of input and m^6^A-eCLIP reads at the 3′ end of the *MGMT* coding sequence in untreated and STM2457-treated U87-R samples. Reproducible U87-R m^6^A-eCLIP sites and DRACH motifs are labeled. (C) m^6^A-IP/qPCR for *MGMT* in U87-R and LN229-R cells. Fold enrichment over input normalized to an unmodified spike-in control RNA is shown. (D and E) The effect of 96 h, 50 μM STM2457 treatment on *MGMT* expression determined by RT-qPCR (normalized to *ACTB*; D) and western blotting (E). (F) Percent of remaining EU-labeled *MGMT* mRNA at increasing times since EU removal in U87-R cells pre-treated for 96 h with 50 μM STM2457 or DMSO, quantified by RT-qPCR (*n* = 3 technical replicates). (G and H) Effect of 72h *IGF2BP2* siRNA knockdown in U87-R cells on *MGMT* mRNA (G) and protein (H) level. (G) Fold change is relative to matched siControl replicate and normalized to *GAPDH* (*n* = 6 biological replicates). Data are mean ± s.d. of 3 biological replicates, unless otherwise specified. Two-tailed Student’s *t*-test; *p < 0.05; **p < 0.01; ***p < 0.001; ****p < 0.0001; ns, not significant.

The central role of *MGMT* in mediating GBM TMZ resistance, its confirmed m^6^A modification in resistant cells, and the canonical role of m^6^A as *repressing* gene expression presented a conundrum, leading us to consider an alternative role for m^6^A in *enhancing* its expression. To explore this idea, we tested the effects of STM2457 on *MGMT* expression. We tested multiple STM2457 doses (Figure S4E) and treatment times (Figure S4F), and found that treatment for 96 hours with 50 μM STM2457 reduced *MGMT* expression at the RNA (Figure 4D) and protein (Figure 4E) levels in U87-R, LN229-R, U251-R cells, and T98G cells. In other words, m^6^A methylation enhances MGMT expression, rather than repressing it as is typical of the YTHDF2-associated mRNA decay mechanism [12,13]. The consistent loss of *MGMT* mRNA and protein with STM2457 treatment suggests that m^6^A has a stabilizing effect, which we confirmed using RNA stability assays. Transcription inhibition-based measurements of RNA decay revealed that *MGMT* is a highly stable mRNA (Figure S4G). To avoid the toxicity of long-term transcription inhibition, we used 5-ethynyl uridine (EU) labeling in a pulse-chase format and observed that *MGMT* mRNA is destabilized upon STM2457 pre-treatment compared to DMSO control, with an observed half life of ∼8 hours versus ∼20 hours, respectively (Figure 4F, Figure S4H). These results are partially consistent with recent work in a glioma stem cell (GSC) model, which found that *MGMT* is m^6^A methylated, but did not map the site [49]. METTL3 knock down in this GSC model also reduced *MGMT* levels, but interestingly the high stability of *MGMT* was not observed in this system. Shi *et al.*also attributed *MGMT* m^6^A methylation to increasing levels of METTL3 enzyme over the course of GSC differentiation, while here we observe highly methylated *MGMT* in the absence of any significant expression changes in m^6^A regulatory enzymes.

While numerous examples of m^6^A-mediated decay have been reported, studies in acute myeloid leukemia were the first to demonstrate that a subset of m^6^A-methylated mRNAs can be stabilized by a second family of m^6^A-binding proteins, the IGF2BP family [10,16]. Interestingly, we find predicted binding sites for IGF2BP2 and IGF2BP3 in the *MGMT* 3’UTR [64]. Our RNA-seq analyses suggest that both *IGF2BP2* and *IGF2BP3* are expressed in our TMZ-R cell lines while IGF2BP1 is not (Figure 3A, Table S1). IGF2BP3 protein levels varied more dramatically across cell lines, and dropped significantly in LN229-R cells relative to parental LN229 cells (Figure S4I), so we focused on *IGF2BP2*. We knocked down *IGF2BP2* with siRNA and found that its depletion reduced *MGMT* mRNA and protein levels (Figure 4G-H). Like *MGMT*, *IGF2BP2* mRNA has m^6^A sites with decreased enrichment in STM2457-treatment U87-R (Figure S4J, Table S4) and decreased expression in response to STM2457 treatment (Figure S4K), suggesting both transcripts are regulated similarly by m^6^A in TMZ-R glioma cells. Taken together, our findings suggest that *MGMT* stabilization by m^6^A is mediated at least in part by IGF2BP2, which itself is regulated in a similar manner, constituting a positive feedback mechanism.

### A broader mechanism of m^6^A-mediated stabilization of TMZ-R associated transcripts

Given the complex and interconnected signaling mechanisms that have been previously associated with TMZ resistance, we were curious whether other TMZ resistance-associated transcripts beyond *MGMT* are also stabilized by m^6^A. Overall, gene expression changes caused by STM2457 treatment in parental and TMZ-R cells were strongly correlated (Figure S5A), which is consistent with the similar m^6^A profiles in parental and resistant cells (Figure 3C-E). However, we noticed that STM2457-induced gene expression changes in TMZ-R cells had a slight negative correlation with resistance-induced expression changes in TMZ-R cells relative to their parental counterpart (Figure S5B), and that this is not the case with STM2457-induced gene expression changes in the parental cell lines (Figure S5C). Indeed, genes significantly upregulated in TMZ-R cells were disproportionately downregulated by METTL3 inhibition (Figure 5A), suggesting that genes that were highly upregulated in TMZ-R cells were also stabilized by m^6^A, similar to *MGMT*. In contrast, genes that were significantly downregulated in TMZ-R cells were largely upregulated by METTL3 inhibition (Figure S5D), possibly reflecting m^6^A-mediated mRNA decay.

**Figure 5.**
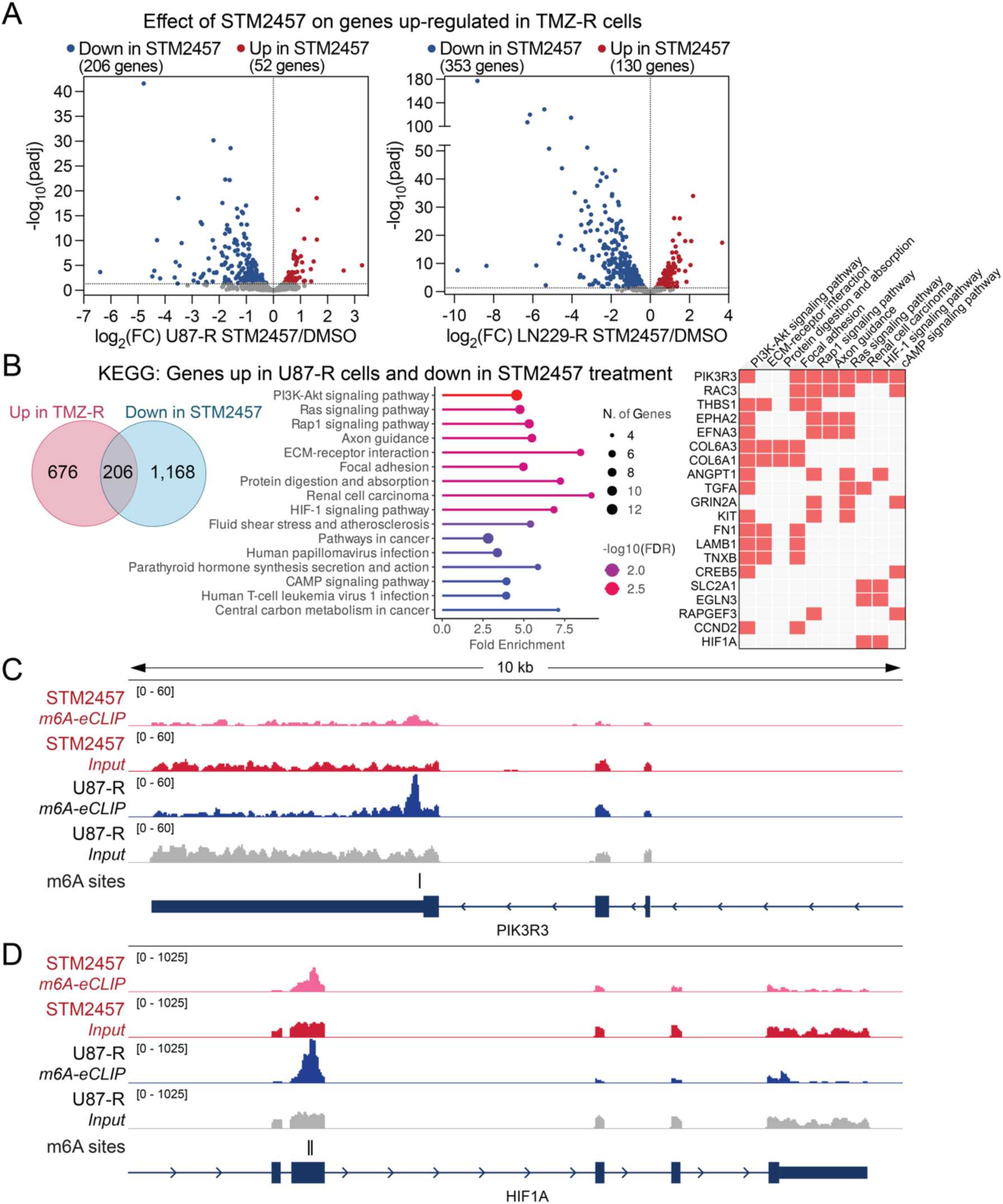
Regulation of TMZ resistance-associated genes by m^6^A. (A) Volcano plots depicting the effect of STM2457 treatment on expression of genes which are significantly upregulated (adjusted p-value < 0.05) in TMZ-R cells. (B) KEGG pathways enriched for genes which are up-regulated in U87-R cells compared to parental U87 MG cells and down-regulated in response to STM2457 treatment in U87-R cells. (C and D) IGV snapshot of input and m^6^A-eCLIP reads at the 3′ ends of *PIK3R3* (C) and *HIF1A* (D) in untreated and STM2457-treated U87-R samples. Loci of single nucleotide m^6^A-eCLIP sites in U87-R samples are labeled.

To explore whether genes that behave like *MGMT* (expression up in TMZ-R relative to the parental line and down with STM2457 treatment) are also associated with TMZ resistance, we looked more closely at the functional annotations associated with this gene set. KEGG pathway analysis revealed that this gene set was enriched for pathways known to be involved in TMZ resistance, including PI3K-Akt signaling, Ras and Rap signaling, extracellular matrix-receptor interactions, and HIF-1 signaling (Figure 5B, Figure S5E). PI3K-Akt signaling is frequently amplified in GBM and contributes to TMZ resistance by promoting cell survival, proliferation, and angiogenesis [65]. PI3K-Akt signaling is tightly interconnected with upstream Ras/Rap signaling. While Ras directly activates PI3K in response to growth factors, Rap proteins also modulate PI3K activity via regulation of focal adhesion and integrin signaling. Additionally, the PI3K-Akt pathway positively regulates hypoxia-inducible factor-1 (HIF-1) signaling and *MGMT* expression, both of which promote cell survival in response to chemotherapy [66–69]. Furthermore, like *MGMT*, many of these key regulatory genes are regulated by m^6^A directly, including PIK3R3 (a regulatory subunit of PI3K) (Figure 5C). Interestingly, m^6^A was previously shown to promote *PIK3R3* mRNA stability via IGF2BP1 and IGF2BP2 in renal cancer, suggesting *MGMT* and *PIK3R3* may be regulated by similar mechanisms [70]. HIF1A, the alpha subunit of the HIF-1 transcription factor, TGFA, a growth factor which activates PI3K signaling, and SLC2A1 (GLUT1), a critical HIF-1 target and proposed GBM therapeutic target, were also among transcripts upregulated in TMZ-R cells and directly stabilized by m^6^A (Figure 5D, Figure S5F). These results suggest that a broader network beyond *MGMT*, including PI3K-Akt signaling and associated pathways, are stabilized by m^6^A methylation to maintain TMZ resistance. Overall, this supports a model in which m^6^A methylation regulates key pathways that are critical for maintaining TMZ resistance, and METTL3 inhibition resensitizes cells to TMZ not only by destabilizing *MGMT*, but by negatively regulating this cohort of mRNAs.

## DISCUSSION

Here we show that m^6^A stabilizes a subset of transcripts associated with acquired TMZ resistance in glioma cell culture models. In particular, we show that *MGMT*, the DNA repair enzyme responsible for repairing TMZ-induced DNA damage, is destabilized by inhibition of the mRNA m^6^A methyltransferase METTL3 by STM2457. While most STM2457-responsive transcripts in these cells demonstrate canonical m^6^A-mediated regulation and are stabilized by METTL3 inhibition, TMZ-R-associated transcripts, like *MGMT*, are destabilized by METTL3 inhibition. Intriguingly, this suggests that while m^6^A-mediated regulation is not dramatically re-wired in these cells, there may be TMZ-R specific mechanisms that alter the functional impact of m^6^A. The identity of TMZ-R specific genes differs across TMZ-R lines, likely reflecting their different basal gene expression patterns. However, *MGMT* is the most highly upregulated gene in all cases and the functions associated with TMZ-R DEGs are shared, including PI3K-Akt signaling and related cellular pathways. Taken together, this suggests that there may be specific features of these genes or feedback mechanisms involving m^6^A reader proteins that underlie this coordinated behavior.

Supporting this idea, we found that transcripts encoding m^6^A regulatory enzymes and binding proteins responded similarly to acquired TMZ resistance and STM2457 treatment compared to their parental line (Figure S4K). Previous work has proposed that METTL3 is upregulated in TMZ-resistant cells, but this is inconsistent with our data showing stable METTL3 expression and bulk level m^6^A methylation [49]. Instead, this points to a small subset of transcripts that contain altered m^6^A sites, which is also consistent with our RNA-seq data. One possible explanation for this is that the stabilizing effect of m^6^A increases the proportion of methylated transcripts of these TMZ-R mRNAs. But the question remains, why are TMZ-R transcripts stabilized while the majority of mRNA is destabilized by m^6^A? The majority of m^6^A binding proteins are also unchanged between parental and TMZ-R cell lines. But intriguingly, the relative expression of m^6^A-binding proteins does seem to change in response to STM2457 treatment, with the levels of YTHDF1 and YTHDF2 increasing, and IGF2BP2 and IGF2BP3 decreasing (Figure S4K). Overall, these STM2457-induced changes would result not only in altered m^6^A status of TMZ resistance-related mRNAs such as *MGMT*, but also change the functional consequences of remaining m^6^A sites via the changes in relative levels of m^6^A-binding proteins.

Beyond the effects of m^6^A-binding proteins, recent work has revealed that interactions between mRNA m^6^A sites and translating ribosomes can also regulate mRNA stability [28–30]. Importantly, these studies revealed that the precise location of the m^6^A site near a stop codon, specifically whether it is in the coding sequence or in the 3’UTR, may be a factor in whether m^6^A stabilizes or destabilizes mRNAs. We find *MGMT* m^6^A-eCLIP peaks spanning both the coding sequence and 3′UTR, but the two *MGMT* m^6^A sites were called in the 3′ end of the coding sequence, with one occurring just upstream of the stop codon. This location of m^6^A adjacent to the stop codon strongly suggests that active translation of m^6^A-methylated *MGMT* may also influence whether m^6^A has a stabilizing or destabilizing effect, but the involvement of this mechanism requires more detailed future study.

Our finding that m^6^A-mediated feedback mechanisms regulate the development and maintenance of TMZ resistance could also have clinical implications. Though alkylation agents are now less commonly used in the clinic, TMZ remains standard of care treatment for GBM patients. *MGMT* promoter methylation is used as a proxy for *MGMT* expression levels, and is a prognostic indicator for patient outcome in response to TMZ [45,46]. Lack of DNA mismatch repair activity has broadly been shown to render patients more resistant to treatment with alkylating agents. Despite this, little is known about *MGMT* expression regulation in response to chronic TMZ treatment. Longitudinal studies in patients have attempted to explore this question but are impeded by limited availability of patient samples at recurrence [71]. A predominant current model is that glioma stem cells drive recurrence because they are more chemoresistant [72], and previous work has compared *MGMT* m^6^A methylation in glioma stem cells relative to differentiated cells [49]. Here we show that *MGMT* status changes in direct response to chronic TMZ treatment, and that the high levels of m^6^A methylation on upregulated *MGMT* transcripts occurs in the background of stable levels of METTL3 expression.

Because *MGMT* is central to reversing TMZ-induced alkylation damage, efforts have been made to target *MGMT* activity and expression as a potential path to re-sensitizing tumors that have become TMZ resistant [43]. *O*^6^-benzylguanine, an irreversible inhibitor of MGMT activity, was tested in Phase I and II clinical trials for its ability to re-sensitize tumors to TMZ but demonstrated high levels of toxicity [47,73]. While alternative approaches have been proposed to re-methylate the *MGMT* promoter and shut down its transcription [74], this is not yet feasible in patients. The data we present here provide a molecular basis for pharmacological inhibition of METTL3 as a potential strategy to sensitize GBM tumors to TMZ; however we note that GBM is currently excluded from ongoing clinical trials of METTL3 inhibitors [37].

Our work also opens many other interesting avenues for future investigation. All of the cell lines used in this study have wild type isocitrate dehydrogenase (IDH) status, but a small proportion of GBM tumors have mutations in these key metabolic enzymes. Specific IDH mutations are known to have neomorphic activity that results in production of the reported oncometabolite 2-hydroxyglutarate [75]. 2-hydroxyglutarate is a known inhibitor of ɑ-ketoglutarate-dependent enzymes, including DNA and RNA demethylases, which could add an additional layer of feedback within the m^6^A regulatory network and influence the effects of METTL3 inhibition. It will also be interesting to dissect at a molecular level how *MGMT* m^6^A sites interact with m^6^A binding proteins and translation machinery, as these events would presumably be mutually exclusive. More broadly, it is not clear how different m^6^A-binding proteins cooperate or compete for binding to a specific m^6^A site, and *MGMT* could provide an interesting case study to investigate this. Finally, our work underscores the value of detailed molecular dissection of m^6^A function on specific transcripts, which can reveal diverse modes of m^6^A-mediated gene expression regulation with meaningful clinical implications.

## METHODS

### Cell culture

U-87 MG, LN-229, and T98G cells were obtained from ATCC and U-251 MG cells were obtained from Sigma-Aldrich. LN-229 cells were cultured in Dulbecco’s Modified Eagle’s Medium supplemented with 5% fetal bovine serum, 1% penicillin/streptomycin, and 1 mM sodium pyruvate (Gibco). U-87 MG, U-251 MG, and T98G were cultured in Eagle’s Minimum Essential Medium (ATCC) supplemented with 10% fetal bovine serum and 1% penicillin/streptomycin. All cell lines were cultured at 37°C and 5% CO2 in a humidified atmosphere. TMZ-R cell lines were generated by continuous maintenance in media supplemented with 50 μM TMZ (Sigma-Aldrich) for at least four weeks.

### Cell viability assays

To measure the effects of TMZ or STM2457 on cell viability, the CyQUANT Cell Proliferation Assay (Invitrogen) was used according to the manufacturer’s specifications and fluorescence was measured using a SpectraMax iD3 microplate reader (Molecular Devices). Cells were seeded 2,500 cells per well in 96-well culture plates and treated the following day by replacing the culture media with media containing drug. For TMZ, cells were treated with 50 μM for 72 h. For STM2457, cells were treated for 96 h with concentrations ranging from 0 to 200 μM. A nonlinear regression, normalized response, variable slope, dose-response model was used to determine the IC50. DMSO concentrations were normalized across all treatment conditions.

Individual experiments were performed with six technical replicates per condition.

### Colony formation assays

For all cell lines, 1,000 cells per well were seeded in 6-well plates and cultured with media containing TMZ (0 or 50 μM) or STM2457 (0, 10, 25, and 50 μM) for 10 days (T98G) or 14 days (U-87 MG, LN-229, and U-251 MG lines). DMSO concentrations were normalized across all treatment conditions. At the end of observation, cells washed with PBS, fixed and stained in 1 mL per well 0.5% crystal violet; 25% methanol for 20 mins with light shaking, then rinsed with MilliQ water.

### Growth curve

U-87 MG parental and TMZ-R cells were cultured in the presence of 50 μM TMZ, 50 μM STM2457, 50 μM TMZ and 50 μM STM2457 combined, or DMSO alone (control). DMSO concentrations were normalized across all treatment conditions. Cells were counted and 1×10^6^ cells were replated in 10-cm dishes with fresh drug every three days and population doublings over the indicated time course were plotted.

### RNA extraction and quantitative real-time PCR

For RT-qPCR and RNA-seq experiments, total RNA was extracted from cells using Trizol (Invitrogen) according to the manufacturer’s instructions. RNA concentration was determined using the NanoDrop 8000 Spectrophotometer (Thermo Scientific). 0.5 μg of total RNA was reverse transcribed using iScript Reverse Transcription Supermix for RT-qPCR (Bio-Rad) according to the manufacturer’s instructions. Real-time PCR was performed using the Bio-Rad CFX96 Real-Time PCR System (Bio-Rad) in 96-well PCR plates using 10 μL reactions composed of 5 μL SsoAdvanced Universal SYBR Green Supermix (Bio-Rad), 1 μL of 4-fold diluted cDNA per reaction, and 200 nM of each forward/reverse primer. All qPCR reactions were performed in triplicate for each sample and analyzed using the 2^-ΔΔCT^ method and normalized to *GAPDH* or *ACTB*, as specified. All steps were performed using qPCR grade water (Fisher Scientific). Primer sequences are provided in Table S5.

### RNA-sequencing

For RNA-seq from U-87 MG cells, RNA was first polyA selected using the Dynabeads mRNA DIRECT Purification kit (Invitrogen), followed by fragmentation using RNA Fragmentation Reagents (Invitrogen) and cDNA library preparation performed using TruSeq Stranded mRNA Library Prep (Illumina). Paired-end 100 bp sequencing was performed on a NovaSeq X Plus instrument (Illumina) with 50 million read pairs. For RNA-seq from LN229 cells, RNA was polyA selected and libraries prepared using the Watchmaker mRNA Library Prep kit (Watchmaker Genomics). Paired-end 100 bp sequencing was performed on a NovaSeq X plus instrument (Illumina) with 25 million read pairs. For all RNA-seq analysis, reads were trimmed using Trimmomatic (version 0.39) [76], aligned to GRCh38 using STAR (version 2.7.11a) [77], and counts determined using Salmon (version 1.4.0) [78].

Differential expression analysis was performed using DESeq2 [79]. An adjusted p-value of 0.05 was considered significant. All RNA-seq experiments were performed with three biological replicates for each condition. Pathway analysis was performed using ShinyGO 0.82 [80] and Enrichr [81–83], and the functional protein association network was generated using STRING v12.0 [84].

### Quantification of DNA methylation

DNA methylation was detected and quantified using the OneStep PLUS qMethyl PCR Kit (Zymo) according to the manufacturer’s instructions. Genomic DNA was extracted from 3×10^6^ cells using the DNeasy Blood & Tissue Kit (QIAGEN). 20 ng of genomic DNA was used for each reaction, and real-time PCR parameters matched those specified by the manufacturer except for the annealing temperature, which was 62.5°C. Three biological replicates were used, and each real-time PCR reaction was performed in technical triplicate, and statistical analysis was performed as recommended by the manufacturer.

### m^6^A immunoprecipitation

For m^6^A immunoprecipitation and quantitative PCR (m^6^A-IP/qPCR), total RNA was extracted from cells using Trizol (Introgen) and then polyA selected using the Dynabeads mRNA DIRECT Purification kit (Invitrogen). 4 μg of polyA-selected RNA was subjected to m^6^A-IP using the EpiMark N6-Methyladenosine Enrichment Kit (NEB) and RT-qPCR was performed as described above. For each sample, prior to immunoprecipitation, 4.05 μg of RNA was diluted to 50 ng/μL and 1 μL of the provided m6A(-) control RNA diluted 1:1,000 was spiked into the RNA samples. 50 ng of each sample was set aside and used as input. RT-qPCR results were plotted as fold enrichment in IP over input relative to the m6A(-) control RNA spike-in.

### m^6^A-eCLIP

For m^6^A enhanced crosslinking immunoprecipitation (m^6^A-eCLIP), 3 biological replicates of U-87 MG parental and TMZ-R cells and 2 biological replicates of U87-R cells treated with 50 μM STM2457 for 96 h were collected. RNA extraction was performed using the QIAGEN RNeasy Midi Kit (QIAGEN) and m^6^A-eCLIP and bioinformatics analysis was performed by EclipseBio, using their proprietary procedures [85]. UMIs were trimmed (umi_tools, cutadapt) [86,87], filtered for repetitive elements and aligned to hg38 (STAR) [77], and PCR duplicates removed (umi_tools) [86]. Peaks and single nucleotide sites were called using CLIPper [88,89] and PureCLIP [90], respectively.

### Immunoblotting

Cells were collected by scraping into ice-cold PBS and pelleted by centrifugation at 1000*g* for 5 min at 4°C. Cell lysis was performed by resuspending cell pellets in three pellet volumes of lysis buffer (50 mM Tris pH 7.4, 300 mM NaCl, 1% NP-40, 0.25% deoxycholate, 1 mM DTT, and 1x SigmaFast protease inhibitor [Sigma-Aldrich]), incubating on ice for 30 min, and clarifying by centrifugation at 20,000*g* for 20 min at 4°C. Protein concentrations determined by Pierce BCA protein assay. 10 μg of each sample were loaded onto 4-12% NuPAGE Bis-Tris gradient gels (Invitrogen) and run at 200V for 50 min in 1X MOPS buffer and then transferred to nitrocellulose membranes (Bio-Rad) by semidry transfer at 17V for 45 min with 2x NuPage Transfer Buffer with 10% methanol. Membranes were blocked for one hour in 5% nonfat dry milk powder resuspended in Tris-Buffered Saline with 0.1% Tween-20 (TBST). Primary antibodies were diluted in 5% nonfat dry milk in TBST and incubated with rocking overnight at 4°C. Antibodies for MGMT (Cell Signalling Technology, 2739), METTL3 (Abcam, ab195352), IGF2BP2 (Abcam, ab124930), and IGF2BP3 (Abcam, ab177477) were used at a 1:1,000 dilution. Anti-GAPDH (Cell Signalling Technology, 5174) served as a loading control and was used at a 1:10,000 dilution. Following incubation with primary antibody, membranes were incubated in donkey anti-Rabbit IgG secondary antibody DyLight 800 (Invitrogen) at a 1:5,000 dilution at room temperature for 2 h with rocking and then imaged using the ChemiDoc MP Imaging System (Bio-Rad).

## LC-MS/MS

Total RNA was extracted from cells using Trizol (Invitrogen) according to the manufacturer’s instructions. mRNA was enriched by polyA selection using the Dynabeads mRNA DIRECT Purification kit (Invitrogen), followed by ribosomal RNA depletion with the NEBNext rRNA Depletion Kit v2 (NEB) and size selection for transcripts greater than 200 nucleotides long with the RNA Clean & Concentrator-5 kit (Zymo). mRNA enrichment was validated by electrophoresis on a Agilent 2100 Bioanalyzer using the RNA 6000 Pico kit (Agilent). The RNA was then digested and analyzed using minor modifications to previously established procedures [91,92]. Specifically, 100 ng of RNA was digested into nucleosides in digestion buffer (50 mM Tris-HCl pH 8 [Sigma], 1 mM MgCl2 [Sigma], 0.2 U/μL bezonase [Thomas Scientific], 0.002 U/μL phosphodiesterase I [Sigma], 0.02 U/μL alkaline phosphatase [Sigma]) at 37°C for 3 h. Samples and standard mixes prepared from pure nucleoside standards were filtered by centrifugation through Millex PVDF sterile syringe filters (0.22 μm, Millipore Sigma) at 10,000 x *g* for 1 min, transferred to autosampler vials (Thermo Scientific), and injected into a Shim-pack GIST C18 (2 μm, 2.1 × 50 mm) reverse phase HPLC column coupled to a Shimadzu 8050 NX Triple Quadrupole mass spectrophotometer. Liquid chromatography was performed using a linear gradient with 0.1% aqueous formic acid to 50% methanol with 0.1% formic acid over 8 min at 400 μL/min. Nucleosides were quantified using retention time and nucleoside to base ion mass transitions, compared to the standard curve obtained from pure nucleoside standards (282.3 > 150.15 for m^6^A; 268.3 > 136.25 for adenosine). A blank containing only digestion buffer was used to control for nucleoside contamination. The m^6^A level was calculated as the percent of m^6^A over adenosine averaged across three injections.

### RNA stability assays

RNA decay over time was measured using the Click-iT Nascent RNA Capture Kit (Invitrogen) according to the manufacturer’s instructions. 2×10^5^ U87-R cells were plated in 35-mm dishes and treated the next day by replacing the media with fresh media containing 50 μM STM2457 or DMSO. After 96 h of STM2457 treatment, cells were incubated with 0.5 mM 5-ethynyl uridine (EU) for 1 hour. Media containing EU was then aspirated, cells were rinsed with fresh media, then grown in fresh media until being collected in Trizol at the indicated time points following EU removal. RNA was extracted using Trizol and subsequent capture of EU-labeled RNA and RT-qPCR was performed according to the manufacturer’s recommendations. Biotinylation of EU-labeled RNA by click reaction was performed using 1 μg EU-labeled RNA and 0.5 mM biotin azide. 0.5 μg biotinylated RNA was captured following RNA precipitation using 50 μL of provided Dynabeads MyOne Streptavidin T1 magnetic beads. For RT-qPCR, cDNA was synthesized using the SuperScript VILO cDNA synthesis kit (Invitrogen) using the RNA captured on the beads as a template and 1 μg of undiluted cDNA was used for qPCR, which was performed as described above. RNA stability assay using Actinomycin D (ActD) was performed by seeding 6×10^5^ U87-R cells in 35-mm dishes and treating the following day with 5 μg/mL ActD.

### siRNA transfection

3.5×10^5^ U87-R cells were reverse transfected in 6-well plates with 25 pmol of ON-TARGETplus human *IGF2BP2* SMARTpool siRNA (L-017705-00-0005) or ON-TARGETplus non-targeting control pool siRNA (Horizon Discovery) using Lipofectamine RNAiMAX Transfection Reagent (Invitrogen) according to manufacturer instructions. After 72 h, cells were collected and the effect of *IGF2BP2* knockdown on MGMT expression was assessed by immunoblotting and RT-qPCR.

## DATA AVAILABILITY

Raw and processed data files from all high throughput sequencing experiments have been deposited in the NCBI Gene Expression Omnibus and will be released on or before publication.

## ACKNOWLEDGMENTS

We would like to thank all members of the Nachtergaele lab for valuable discussion and feedback, Olivia S. Rissland for feedback on the manuscript, and Fabian Menges and the Yale Chemical and Biophysical Instrumentation Center for mass spectrometry support. We would also like to thank the

Yale Centers for Genome Analysis and Research Computing for support with RNA sequencing and analysis. This work was supported by a Distinguished Scientist Award from the Sontag Foundation, an NIGMS MIRA award (R35GM146919), and NHGRI R01HG011868 to S.N. E.D. was previously supported by training grants T32GM007499 and T32GM145469, and is currently supported by an NIH NRSA F31 fellowship (F31CA306101). R.P. was supported by an award from the American Cancer Society Center for Innovation in Cancer Research Training. Research reported in this publication was also supported by the National Institute of General Medical Sciences of the National Institutes of Health under award number 1S10OD030363-01A1 (Yale Center for Genome Analysis).

## Supplemental Information

**Figure S1.**
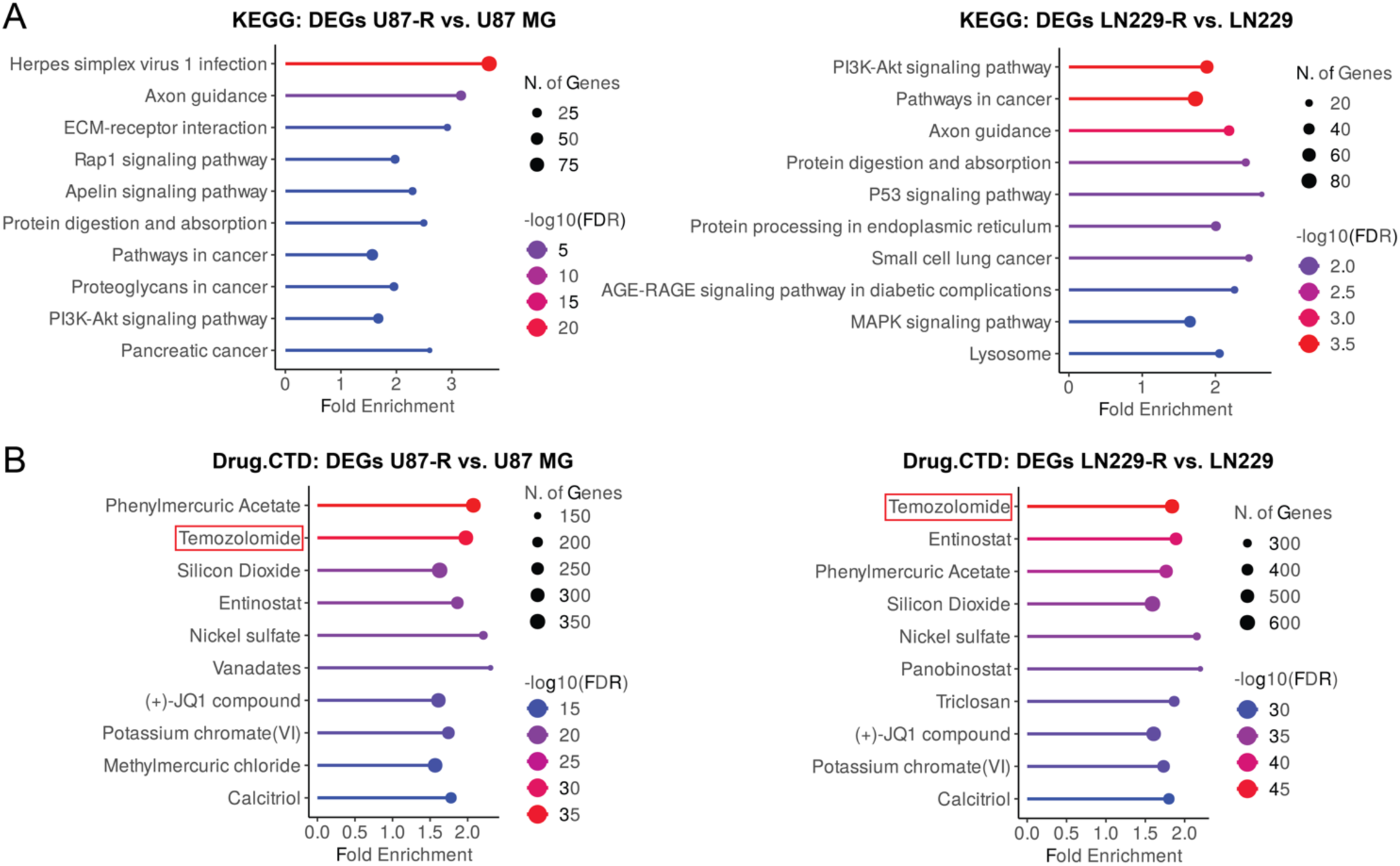
Gene set enrichment analyses for DEGS IN TMZ-R and parental cell lines, related to. **Figure 1**. (A) Top ten most significantly enriched KEGG pathways by false discovery rate (FDR) for genes differentially expressed (DEGs) between TMZ-R and parental cells. (B) Enrichment of DEGs in TMZ-R vs. parental cells for drugs using the Comparative Toxicogenomics Database (CTD).

**Figure S2.**
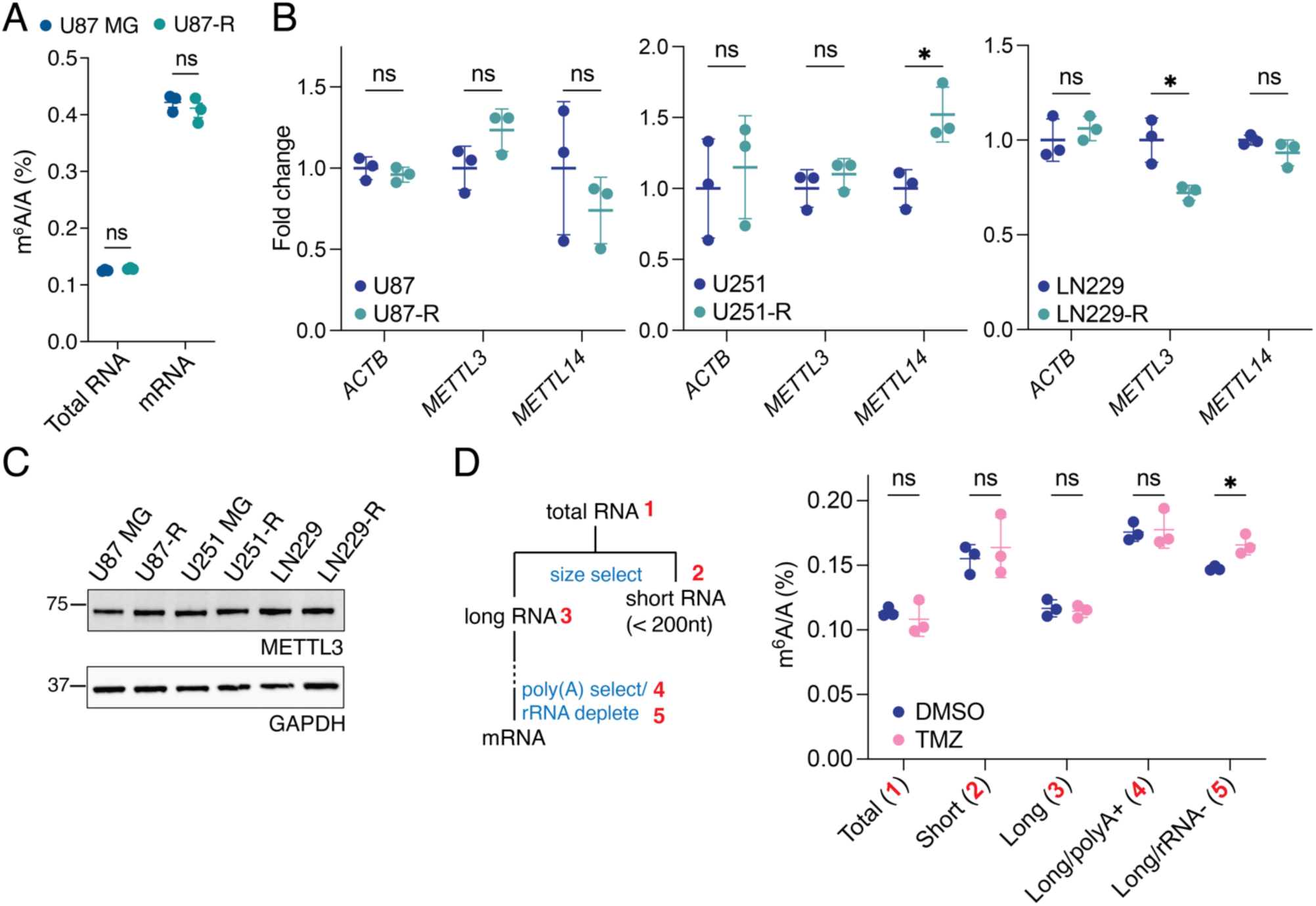
Comparison of bulk m^6^A levels in TMZ-R and parental cell lines, related to. **figure 2**. (A) LC-MS/MS quantification of m^6^A levels in total RNA and mRNA from U87 MG parental and TMZ-R cells expressed as percent m^6^A over unmodified adenosine in total RNA. (B) RT-qPCR comparing expression of m^6^A methyltransferases *METTL3* and *METTL14* between parental and TMZ-R cells. The housekeeping gene *ACTB* served as a negative control. (C) Western blot analysis of METTL3 in parental and TMZ-R cells. (D) LC-MS/MS quantification of m^6^A levels of various RNA fractions collected from U87 MG cells after treatment with 100 μM TMZ for 72 h (*n* = 3 injections). Data are mean ± s.d. of 3 biological replicates, unless otherwise specified. Two-tailed Student’s *t*-test; *p < 0.05; ns, not significant.

**Figure S3.**
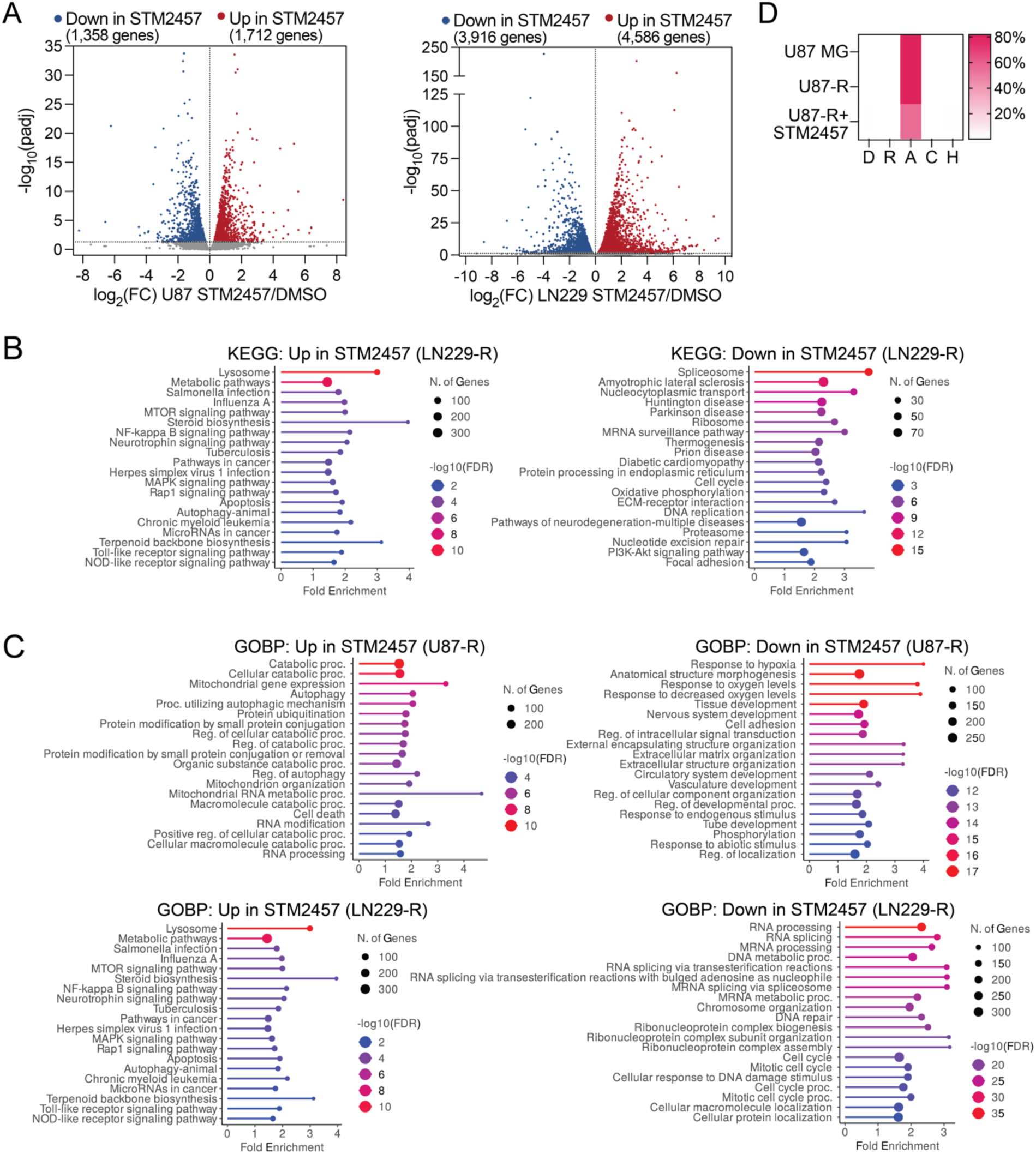
Effect of STM2457 treatment on gene expression, related to. **Figure 3**. (A) Volcano plots showing differentially expressed genes in parental U87 MG or LN229 cells treated with 50 μM STM2457 for 96 h. (B and C) KEGG pathway (B) or Gene Ontology Biological Process (GOBP; C) analysis for genes differentially expressed in response to STM2457 treatment in TMZ-resistant glioma cell lines. (D) Heat map showing the percent of reproducible m^6^A-eCLIP sites in each condition mapped to each position within the DRACH motif. Nucleotides are defined using IUPAC codes (D = G/A/U; R = G/A; H = U/A/C).

**Figure S4.**
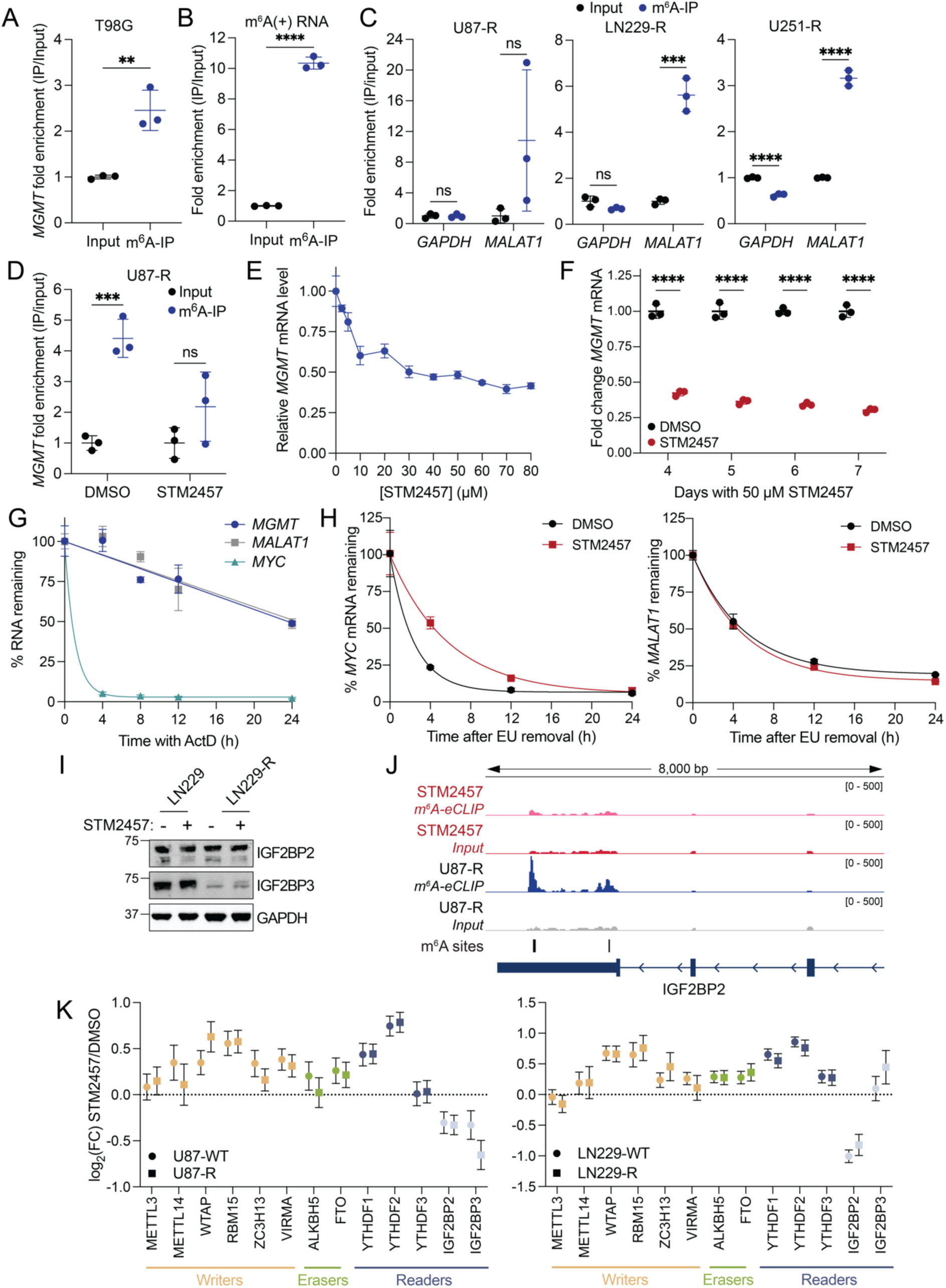
Validation of regulation of *MGMT* by m^6^A, related to. **Figure 4**. (A) m^6^A-IP/qPCR for *MGMT* in T98G cells. (B and C) m^6^A-IP/qPCR in T98G, U87-R, and LN229-R, and control m^6^A-IP for spike-in m^6^A modified RNA (m^6^A(+)). (D) m^6^A-IP/qPCR for *MGMT* in U87-R cells treated with DMSO or 50 μM STM2457 for 96 h. For (A-D), fold enrichment over input normalized to an unmodified spike-in control RNA is shown. Data from U87-R and LN229-R represents 3 biological replicates, while data from T98G and control IP represents 3 technical replicates. (E and F) RT-qPCR measuring *MGMT* expression in U87-R cells after treatment with increasing concentrations of STM2457 for 72 h (E) or treatment with 50 μM STM2457 at increasing incubation times (F). Relative *MGMT* mRNA levels are normalized to *ACTB*. (G) Percent of RNA remaining in U87-R cells after increasing incubation times with the transcription inhibitor Actinomycin D (ActD), quantified by RT-qPCR. (H) Percent of remaining EU-labeled RNA at increasing time points after EU removal, quantified by RT-qPCR. Data are mean ± s.d.. Two-tailed Student’s *t*-test; **p < 0.01; ***p < 0.001; ****p < 0.0001; ns, not significant. (I) Western blot showing IGF2BP2 and IGF2BP3 protein expression in LN229 and LN229-R cells treated with 50 μM STM2457 for 96 h or DMSO. (J) Genome browser snapshot of input and m^6^A-eCLIP reads at the 3′ end of *IGF2BP2* in untreated and STM2457-treated U87-R samples. Reproducible U87-R m^6^A-eCLIP sites are labeled. (K) The effect of STM2457 treatment on expression of m^6^A regulators, determined by RNA-seq. Error bars represent standard error of log2(fold change) (*n* = 3).

**Figure S5.**
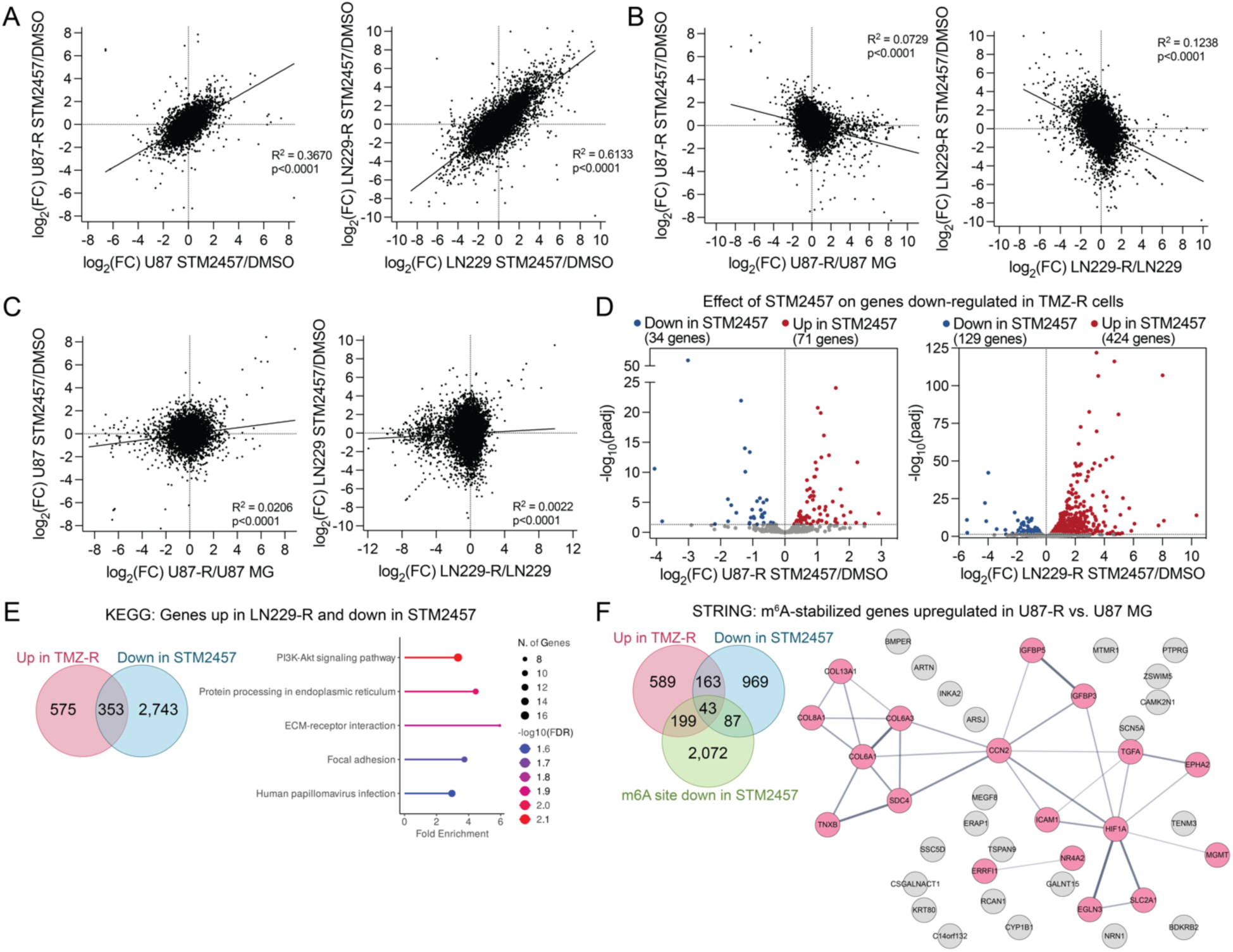
Relationship between acquired TMZ resistance and regulation of gene expression by m^6^A, related to. **figure 5**. (A) Linear regression comparing the effect of METTL3 inhibition (STM2457- versus DMSO-treatment) on gene expression in parental and TMZ-R glioma cells. (B-C) Linear regression comparing the effect of TMZ resistance (TMZ-R vs. parental cells) and METTL3 inhibition (STM2457 vs. DMSO-treatment) in TMZ-resistant (B) and parental (C) U87 MG and LN229 on gene expression. (D) Volcano plots depicting the effect of STM2457 treatment on expression of genes which are significantly downregulated in TMZ-R cells. (E) KEGG pathways enriched for genes which are up-regulated in LN229-R cells compared to parental LN229 and down-regulated in response to STM2457 treatment in LN229-R cells. (F) STRING functional protein association network (interaction score > 0.4) for genes which are significantly up-regulated in U87-R cells (compared to parental U87 MG), down-regulated by STM2457 treatment, and have a m^6^A-eCLIP site in U87-R with decreased enrichment following STM2457 treatment. Node thickness indicates the strength of data support.

